# Single-molecule tracking reveals the dynamics of Ipl1 recruitment to the kinetochores and spindles in *S. cerevisiae*

**DOI:** 10.1101/2024.08.01.606162

**Authors:** Nitesh Kumar Podh, Ayan Das, Akriti Kumari, Kirti Garg, Rashmi Yadav, Kirti Kashyap, Sahil Islam, Anupam Gupta, Gunjan Mehta

**Affiliations:** Laboratory of Chromosome Dynamics and Gene Regulation, Department of Biotechnology, Indian Institute of Technology Hyderabad, India; Single-Molecule Science, University of New South Wales, Sydney, Australia; Department of Physics, Indian Institute of Technology Hyderabad, India

## Abstract

Aurora kinase B, Ipl1 in *Saccharomyces cerevisiae*, is the master regulator of cell division required for checkpoint regulation, spindle assembly and disassembly, chromosome segregation, and cytokinesis. Decades of research employed ensemble averaging methods to understand its dynamics and function; however, the dynamic information was lost due to population-based averaging. Here, we use single-molecule imaging and tracking (SMIT) to quantify the recruitment dynamics of Ipl1 at the kinetochores and spindles in live cells. Our data suggest that Ipl1 is recruited to these locations with different dynamics. We have demonstrated how the recruitment dynamics of Ipl1 at the kinetochores during metaphase changes in the presence and absence of tension across the kinetochore, in the absence of protein phosphatase 1 (Glc7), and the absence of its known recruiters (Ctf19 and Bub1). The SMIT of other chromosome passenger complex members suggests its hierarchical assembly at the kinetochore. Hence, SMIT provides a dynamic view of the Ipl1 trafficking at the kinetochores and spindles.

## INTRODUCTION

Aurora kinases (AKs) play essential roles during cell division (mitosis and meiosis). They spatiotemporally phosphorylate various proteins at the kinetochores and spindles for functions such as kinetochore assembly, checkpoint regulation, spindle assembly/disassembly, chromosome segregation, and cytokinesis (Figure 1A) (1). The impaired function of AKs leads to severe chromosome missegregation, aneuploidy, and cancers. The human genome contains three genes encoding AKs: AK-A, AK-B, and AK-C. Their overlapping localization, functions, and substrates make studying specific functions/dynamics of AK-B in isolation challenging. In yeast *S. cerevisiae*, Ipl1 (Increase-in-ploidy) is a single essential AK, homologous to human AK-B, that serves as a valuable tool for studying its dynamics in isolation. Like human AK-B, Ipl1 localizes to the centromeres/kinetochores (during metaphase for kinetochore assembly and spindle checkpoint functions). It relocates to the spindles and spindle midzones (during anaphase and late anaphase for spindle assembly and disassembly, respectively, Figure 1A) (2, 3). Ipl1 is recruited to the kinetochores or spindles as a part of the Chromosome Passenger Complex (CPC, made up of Survivin (Bir1), Borealin (Nbl1), and INCENP (Sli15)). Sli15 links the Ipl1 and the Bir1-Nbl1 complex (Figure 1B). Two pathways are known to recruit the CPC complex to the kinetochores: 1) Bub1/Sgo1 mediated pathway that tethers the CPC to the inner kinetochore protein Ndc10 (4–7), 2) Sli15 mediated targeting of the CPC to the inner kinetochore protein Ctf19 (COMA complex) (8–9). CPC is recruited to the spindle by the Microtubule Binding Domain (MTB) of Sli15 (10). Also, recent reports have demonstrated three discrete binding sites of the CPC complex: inner centromere, inner kinetochore, and spindles (11–12). Conversely, two recent studies have demonstrated that the inner centromere and the inner kinetochore CPC targeting mechanisms are at least partially redundant for chromosome biorientation and cell viability in budding yeast (8–9). Additionally, mutations in the SAH domain of Sli15 prevent CPC localization to all three sites in budding yeast (12). All this evidence suggests that Ipl1 localization to all these three sites contributes to substrate phosphorylation at the outer kinetochore (11–13).

**Figure 1.**
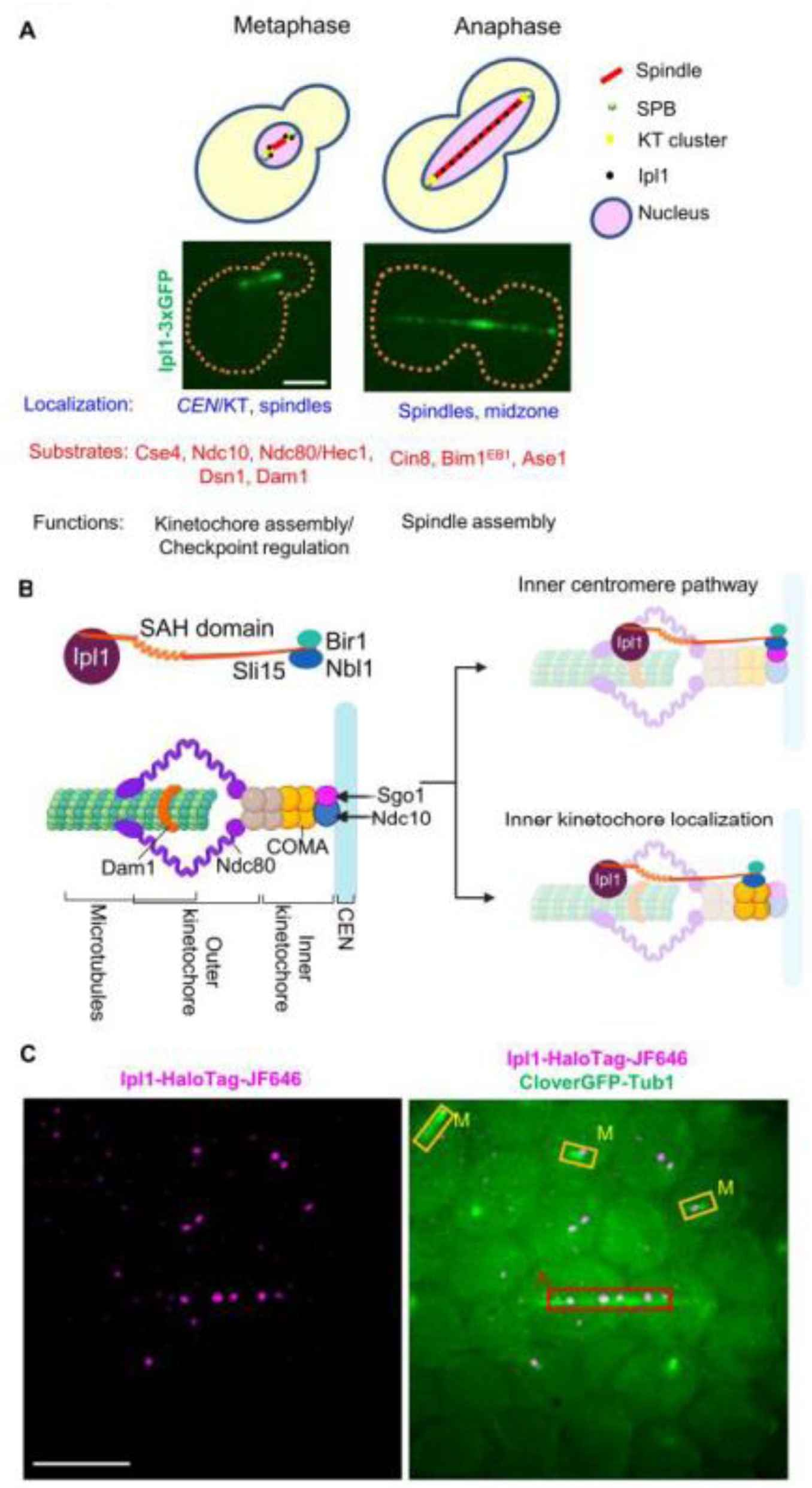
Introductory diagrams and the design of the experiment: **(A)** Schematic representation and corresponding microscopic images for the spatiotemporal localization of Ipl1 during mitosis. Ipl1 localizes to the kinetochores (during metaphase) and spindles (during anaphase). Its substrates and functions are depicted. Scale: 2 µm **(B)** Mechanisms of CPC recruitment to the kinetochores: Composition of the kinetochore and CPC. (Right) Reported sites and pathways of CPC localization at the kinetochore. **(C)** Strategy for tracking Ipl1-HaloTag-JF646 at different stages of mitosis at different locations. The left panel shows single molecules of Ipl1-HaloTag-JF646 (filtered image), and the right panel shows a merged image of CloverGFP-Tub1 with Ipl1-HaloTag-JF646. Based on spindle morphology, cells can be classified into metaphase and anaphase, and ROIs can be drawn to track the molecules within that area. Scale: 5 μm. SPB: Spindle-pole body, *CEN*: centromere, KT: kinetochore.

Glc7 is a protein phosphatase 1 (PP1) in yeast *S. cerevisiae,* and it dephosphorylates the Ipl1 substrates at the kinetochores (14). The interplay between Ipl1 and Glc7 still needs to be clearly understood to maintain the critical phosphorylation levels of Ipl1 substrates. Unlike the kinetochore/spindle-specific localization of Ipl1, Glc7 is localized in the entire nucleus throughout the cell cycle (14). However, its specific localization to the kinetochores/Spindle-Pole Bodies (SPBs) is reported during anaphase (14–15).

Protein phosphorylation and dephosphorylation are dynamic processes in which a single protein molecule switches between ON and OFF phosphorylation states because of the simultaneous presence of kinases and phosphatases in live cells (14–16). Decades of research on AK-B/Ipl1 visualized and quantified its bulk localization in cells (using ChIP, immunofluorescence, western blotting, and live cell imaging) and quantified the phosphorylation level of its substrates after fixation and from the cell population. Hence, the dynamic behavior of individual molecules of Ipl1 and the dynamics of substrate phosphorylation were lost in the bulk measurements. The resulting static picture informs that Ipl1 localizes to several places and phosphorylates several substrates (Figure 1A). However, the dynamic information about Ipl1 action could not be quantified, such as how long Ipl1 binds to the centromeres/kinetochore and spindles for phosphorylating its substrates, how Ipl1 finds its target sites by exploring the entire nuclear space (target-search mechanism), how different modulators alter the recruitment dynamics of Ipl1 at the kinetochores/spindles.

In the last decade, single-molecule imaging and tracking (SMIT) has revolutionized how we understand biological processes (transcription, DNA replication, telomerase action) by revealing the dynamics of cellular processes at the single-molecule level (17–23). However, the dynamic regulation of any kinase/phosphatase is not thoroughly quantified despite their dynamic interplay in regulating various biological signaling cascades. In this report, we use the single-molecule imaging and tracking (SMIT) method to quantify the fast dynamics of Ipl1 in live yeast cells. The Ipl1 shows fast exchange/turnover at the kinetochores compared to the spindles, suggesting the importance of dynamic turnover in regulating kinetochore function (kinetochore assembly/checkpoint regulation). Upon depleting Glc7 during metaphase, the residence time and the fraction of specific bound molecules of Ipl1 increase at the kinetochores, suggesting that the fast exchange of Ipl1 is essential to keep the Glc7 away from its kinetochore substrates. Ipl1 is best known for its role in tension sensing at the kinetochores during metaphase for error correction. We quantified the dynamics of Ipl1 at the kinetochores during metaphase in the presence and absence of tension. Our data suggests that once the biorientation is achieved and the kinetochores are under tension, Ipl1 diminishes its binding at the kinetochores. Upon releasing the tension by depolymerizing microtubules, Ipl1 relocates to the kinetochores. Hence, tension across the kinetochores keeps the Ipl1 away, whereas reduced tension, irrespective of erroneous attachment, recruits Ipl1 to the kinetochores. We tracked the dynamics of Ip1l in the absence of Ctf19 or Bub1, known recruiters of Ipl1 at the kinetochores. Our data suggests that the absence of Ctf19 reduces the specific bound fraction of Ipl1 at the kinetochore and increases the search time of Ipl1 to find the kinetochores, whereas the absence of Bub1 abolishes the specific binding of Ipl1 to the kinetochore. SMIT of the other members of the CPC suggests a hierarchical assembly of it at the kinetochores during metaphase based on their residence times. Nbl1-Bir1 assembles first at the kinetochores, followed by the Sli15-Ipl1 complex. In summary, this study reveals the dynamics of Ipl1 recruitment and its target-search mechanism to achieve its multifaceted functions during cell division.

## MATERIAL AND METHODS

### Genetic engineering of yeast strains for SMIT of Ipl1

*S. cerevisiae* strains used in this study were derived from the haploids BY4741 and BY4742, isogenic to S288C (Research Genetics/Invitrogen, Huntsville, AL) (see Table S1 for genetically engineered yeast strains and Table S2 for plasmids). Standard methods were used for yeast transformation using PCR-based homologous recombination (36). For gene deletion and C-terminal tagging of proteins, appropriate PCR fragments were amplified (see Table S3 for primers) and were integrated into the genome by homologous recombination (37–38). All deletions and C-terminal tags were confirmed by either diagnostic PCR or observation of the localization of the appropriate fusions by fluorescence microscopy. For SMIT experiments, the *PDR5* gene, which codes for the membrane transporter protein Pdr5, was deleted to allow the HTL (JF646-HTL) retention inside the yeast cell (21). To visualize single molecules of Ipl1, the *HaloTag-TRP1* cassette was amplified from pTSK573 and transformed into the yeast cells for endogenous integration at the c-terminus of *IPL1*. As a localization marker, *CloverGFP-TUB1* was integrated at the endogenous *TUB1* locus using a plasmid *pHIS3p:CloverGFP-TUB1+3’UTR::URA3*, digested with BsaB1 to integrate at the *TUB1* locus. For *NDC10-GFP* tagging, we used pTSK405 to amplify the *3xGFP::URA3* cassette for endogenous tagging. For auxin-inducible degradation of Cdc20 and Glc7, the endogenous copies were c-terminally fused with *AID*-6HA-Hyg* (39) by amplifying the cassette from the plasmid *pHyg-AID*-6HA*. *pADH1-AFB2-CYCT1* cassette was integrated at the *TRP1* locus using pTSK559 (digested with SwaI).

### Culturing cells for single-molecule imaging

The cells were streaked on the CSM plate from the glycerol stocks (from the -80 °C freezer) and incubated at 30 °C for 48 hours. A single colony was inoculated in 5 ml CSM broth and grown at 30 °C under shaking conditions (230 RPM) for 20-24 hr. 50 µl of this culture was inoculated in fresh 3 ml CSM broth and grown at 30 °C under shaking conditions (230 RPM) for 5-6 hr to bring the cells to the log phase. 1 ml cell suspension was taken in a 14 ml round bottom PP test tube with a snap cap; JF646-HTL was added at 30 nM concentration and kept for shaking for 30 min. Cells were pelleted by centrifugation (2000 RPM for 2 min) and washed twice with 3 ml of fresh prewarmed CSM media to remove unbound JF646-HTL (this step is optional, as unbound JF646-HTL will not give fluorescence). Cells were finally resuspended in 20 µl of CSM media. 3 µl of this cell suspension was taken on the LabTekII imaging chamber, covered by a nutrient agarose pad (8 x 8 mm, CSM + 2% SeaKem GTG agarose), and the cells were then imaged for ∼1 hr using a single-molecule imaging microscope. Every hour, fresh cells were taken for imaging.

To arrest the cells at metaphase (using *CDC20-AID\**), 1 mM auxin was added to the 5-6 hr grown log phase cells and kept for shaking for an additional 2 hr, followed by incubation with JF646-HTL (30 nM) for a further 30 min (Figure S2A). Cells were pelleted by centrifugation (2000 RPM for 2 min) and washed twice with 3 ml of fresh prewarmed CSM + 1 mM auxin media to remove unbound JF646-HTL. Cells were finally resuspended in 20 µl of CSM + 1 mM auxin media. 3 µl of this cell suspension was taken on the LabTekII imaging chamber, covered by a nutrient agarose pad (8 x 8 mm, CSM + 1 mM auxin + 2% SeaKem GTG agarose), and the cells were then imaged for ∼1 hr.

To simulate the condition for kinetochores without tension (Figure 3B, right panel), benomyl treatment (90 µg/µl) for 1 hr (including the last 30 min incubation with 30 nM JF646-HTL) was given after 2 hrs of auxin treatment (Figure S2A). Cells were pelleted by centrifugation (2000 RPM for 2 min) and washed twice with 3 ml of fresh prewarmed CSM + 1 mM auxin + 90 µg/µl benomyl to remove unbound JF646-HTL. Cells were finally resuspended in 20 µl of CSM + 1 mM auxin + 90 µg/µl benomyl media. 3 µl of this cell suspension was taken on the LabTekII imaging chamber, covered by a nutrient agarose pad (8 x 8 mm, CSM + 1 mM auxin + 90 µg/µl benomyl + 2 % SeaKem GTG agarose), and the cells were then imaged for ∼1 hr.

**Figure 2:**
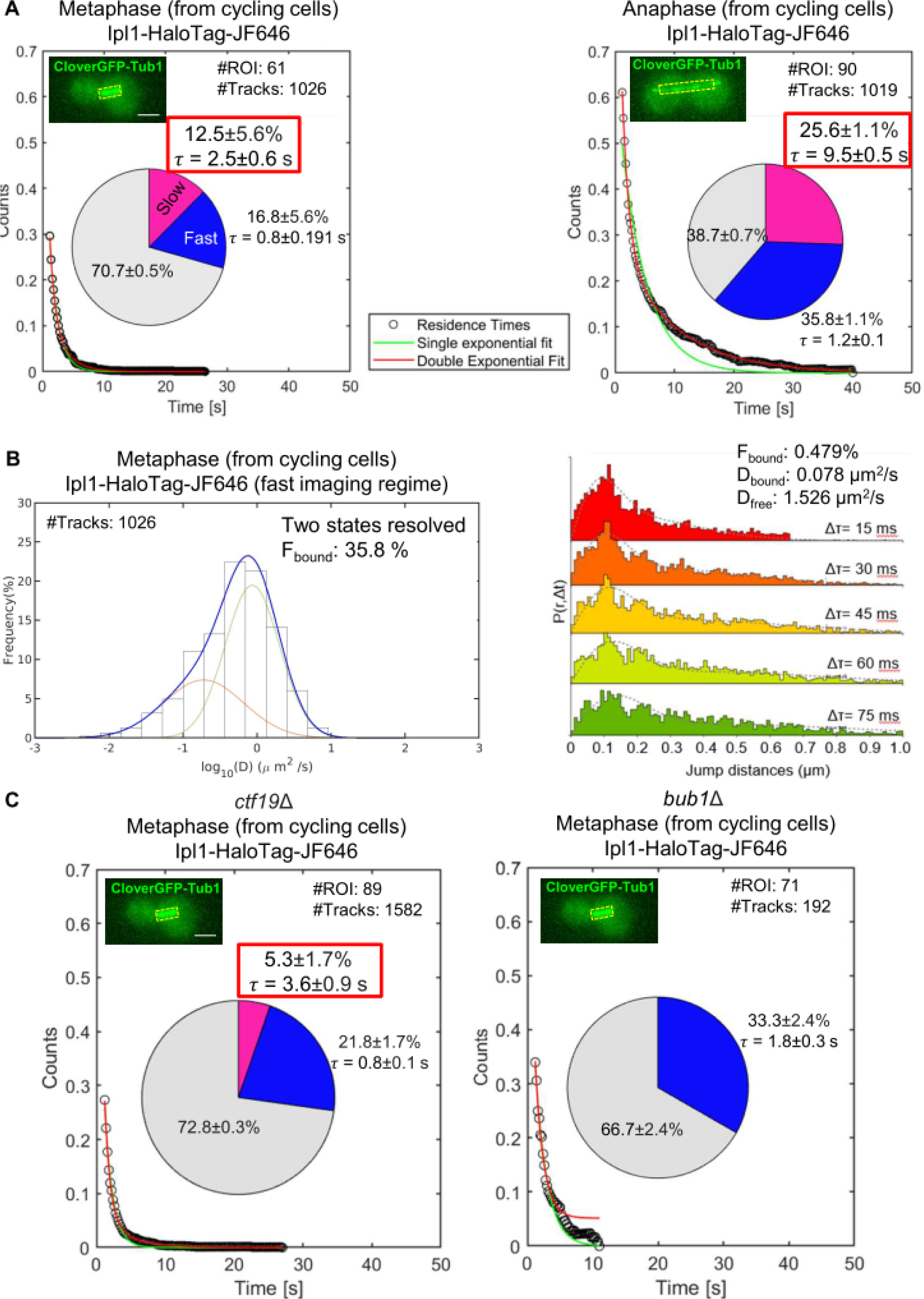
SMIT of Ipl1-HaloTag-JF646 at different stages of mitosis and in the absence of Ctf19 and Bub1: **(A)** Residence time analysis of Ipl1 during different stages of mitosis using a slow imaging regime. Survival probability distributions were fitted with single (green)-or double (red)-exponential decay curves. For all stages, the survival distribution fitted best with the double exponential decay, suggesting two populations of residence times. The pie charts represent the fraction of molecules bound with long residence time (pink fraction) and the fraction of molecules bound with short residence time (blue fraction), along with their mean residence times (τ). The grey fraction represents diffusing molecules. “#Tracks” represents the total number of tracks analyzed, and “#ROIs” represents the total number of cells tracked. The inset images show the ROIs used for tracking. Scale: 2 µm **(B)** SMIT of Ipl1-HaloTag-JF646 at metaphase using a fast-imaging regime. A Histogram of LogD was used to identify the number of states of the molecules. Spot-On based kinetic modelling was used for the robust quantification of the mean D value of the bound and free molecules. **(C)** SMIT of Ipl1-HaloTag-JF646 in the absence of Ctf19 and Bub1 during metaphase using a slow imaging regime. **S**urvival probability distributions and pie charts are represented as Figure 2A. “#Tracks” represents the total number of tracks analyzed, and “#ROIs” represents the total number of cells tracked. The inset images show the ROIs used for tracking. Scale: 2 µm

**Figure 3:**
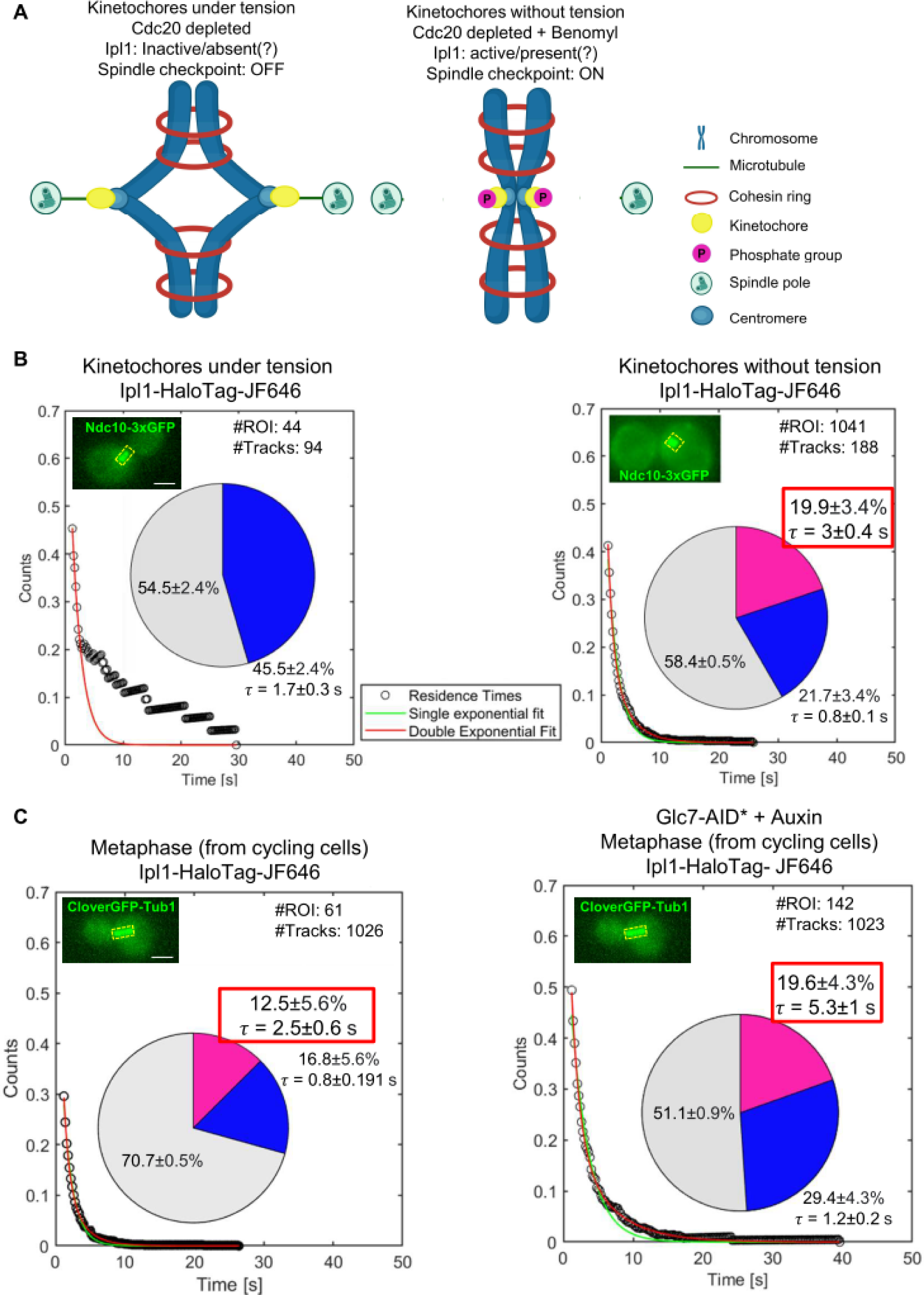
Effect of tension across the kinetochores and absence of Glc7 on the dynamics of Ipl1 at metaphase: **(A)** A schematic representation of the simulated experimental conditions for keeping the kinetochores under tension and releasing the tension. By depleting Cdc20, cells were arrested at metaphase with kinetochores under tension due to the opposite pulling forces exerted by the spindles. The addition of benomyl relieves tension across the sister kinetochores. **(B)** SMIT of Ipl1-HaloTag-JF646 in the presence and absence of tension at the kinetochores during metaphase. Survival probability distributions and pie charts are represented as Figure 2A. “#Tracks” represents the total number of tracks analyzed, and “#ROIs” represents the total number of cells tracked. The inset images show the ROIs used for tracking. Scale: 2 µm **(C)** SMIT of Ipl1-HaloTag-JF646 in the presence and absence of Glc7 at metaphase. Survival probability distributions and pie charts are represented as Figure 2A. The data shown on the left panel is the same as it is shown in Figure 2A (for metaphase), however, it has been shown again for easy comparison with the absence of Glc7. “#Tracks” represents the total number of tracks analyzed, and “#ROIs” represents the total number of cells tracked. The inset images show the ROIs used for tracking. Scale: 2 µm

### Single-molecule imaging

Single-molecule imaging was performed on a Leica DMi8 infinity TIRF inverted fluorescence microscope equipped with a 100X 1.47 NA oil immersion objective lens, a Photometric Prime95B sCMOS camera, and a 638 nm 150 mW laser (referred to as “single-molecule imaging microscope.” The microscope, camera, and lasers were controlled by the Leica LasX software version 3.8.6. Cells were focused using the FITC channel under the wide field illumination, and the best field was selected based on the fluorescence from CloverGFP-Tub1 or Ndc10-3xGFP. Single-focal plane time-lapse movies were acquired with two imaging regimes: 1) the ‘fast-imaging’ regime acquires time-lapse movies with 15 ms time intervals (10 ms exposure time, 100% laser power) to quantify the diffusive behavior of Ipl1 (Figure S2B). However, extensive photobleaching due to high laser power and continuous imaging curtails the estimation of the residence times. 2) The ‘slow-imaging’ regime acquires time-lapse movies with 200 ms time intervals, including 50 ms exposure time, at 30% laser power (Figure S2B) (24). The low laser power and 200 ms time interval allow the molecules to be visible for a long time before they photobleach. Also, the long exposure (i.e. 50 ms) allows the fast-diffusing molecules to blur out due to motion blur and selectively visualize the chromatin-bound molecules to estimate the residence times. Combining the two imaging regimes provides a holistic and quantitative view of various diffusive behaviors and kinetic subpopulations. Images were processed with ImageJ for brightness and contrast adjustments and merging channels.

### Single-molecule tracking

#### Estimation of Bound molecules *C_eq_*

The particle tracking was performed on the “slow imaging” movies using MATLAB-based “TrackRecord” software (20,24,39,41). The software provides automated features for particle detection (puts a threshold on intensity), tracking (using the nearest neighbour algorithm with molecules allowed to move a maximum of 6 pixels from 1 frame to the next, and only tracks that are at least four frames or longer are kept. Gaps to close 4 frames to compensate for fluorophore blinking), photobleaching correction, and quantification of residence time (survival probability distribution).

Since histone H3 (Hht1) molecules are known to be stably bound to chromatin, we used its trajectory as a reference to define bound molecules for other proteins. A maximum distance moved by the histone H3 molecules is extracted using a histogram of displacement (Figure S2C). We identify binding events for the molecule of interest by analyzing trajectory data and extracting segments where the distance moved between consecutive frames, for about 99% of the molecule, is less than or equal to this maximum displacement, *r_min_* (470 nm) and the end-to-end displacement for each particle is greater than a cutoff, *r_max_* (610 nm).

Also, if a molecule diffuses freely but comes back to a distance less than *r_min_* in consecutive frames, it should not be counted as a bound molecule. Therefore, the molecule has to fulfill a minimum number of consecutive frames (*N_min_*) cutoff to be counted as bound (Figure S2D). The *N_min_* value used for 200 ms time-interval movies was 7. Now, the bound fraction *C_eq_* is estimated by examining all trajectories and calculating the number of frames at which the molecules were classified as bound divided by the total number of frames.

#### Estimation of target-search parameter *τ_search_*

Fitting of the survival probability curve

To extract dwell time, the survival distribution, S(t), i.e., *C_eq_*(1-*CDF*), is fitted using the method of least squares to a double exponential decay:

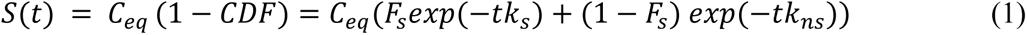

where, *k*_*s*_ and *k*_*ns*_ are the dissociation rates of the molecule to specific and nonspecific binding sites. Considering *τ*_*s*_/*τ*_*ns*_ to be the average dwell time for specific/nonspecific binding, *k*_*s*_ and *k*_*ns*_are inverse of *τ*_*s*_ and *τ*_*ns*_respectively. *F*_*s*_(1 − *F*_*s*_)is the fraction of specific (nonspecific) binding molecules to total bound molecules. CDF is the cumulative distribution function of dwell time.

The bound fraction estimated by the method mentioned above underestimates the true bound fraction as only those particles that are bound for the frames more than *N*_*min*_ are counted. To address this underestimation of bound molecules and thus estimating a “true” bound fraction extrapolation of the survival probability plot back to time t=0 is done. That is why the S(t) curve is fitted to *C*_*eq*_(1 − *CDF*).

To check for overfitting, the distribution is also fit to a single-component exponential:

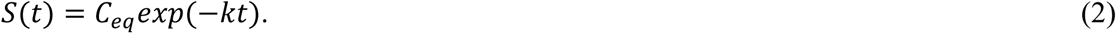

The fits are compared using an F-test to ensure that the two-component model significantly improves over the single-component decay.

### Average dwell time calculation

The average residence time is then estimated as follows:

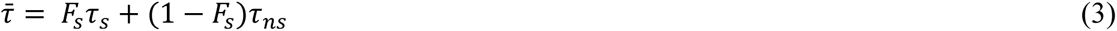

From *F*_*s*_ we can obtain the number of encounters with nonspecific binding sites before encountering a specific binding site, *N*_trails_ = 1/*F*_*s*_ and from *C*_*eq*_ we can get the average free time between two binding events, *τ*_3*D*_ by,

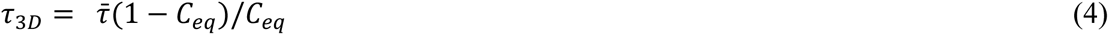

So, the search time, i.e., the time for locating a specific site, can be given as:

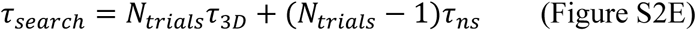

The differences in the residence times or fractions of specific bound molecules between two conditions were statistically checked by the Z-test to derive the p-values.

### For diffusion analysis

For quantifying diffusion parameters (fraction of bound/unbound molecules and mean diffusion coefficient), particle tracking was performed using the “fast imaging” movies by DiaTrack Version 3.05, with the following settings as previously described (19,24). remove blur 0.07, remove dim 45-150, maximum jump 6 pixels, where each pixel was 110 nm. This software determines the precise position of single molecules by Gaussian intensity fitting and assembles particle trajectories over multiple frames. The trajectory data exported from Diatrack was further converged into a single.csv file using a custom computational package, ‘Sojourner.’

We used SPTAnalyzer, the in-house MATLAB script for diffusion analysis. In this analysis, we have discarded tracks with at most six displacements. After filtering the data, we fitted the 30 ms to 75 ms range linearly to determine the diffusion coefficient (D). The D for that specific track was determined by dividing the slope of the linear fitting with R^2^≥0.8 criteria by 4. We have kept the bin-width 0.26 μm. The fitting yields the fractions of the bound (*F_bound_*) and unbound molecules (*F_free_*) by fitting two Gaussians.

Quantifying diffusion parameters using MSD-based logD histograms can produce unreliable results because this method is highly dependent on the choice of track length, and linear fits to MSD may yield misleading diffusion coefficients (D values) for very stable molecules. Therefore, MSD-based analysis can be used to identify resolvable states (e.g., 2 states or 3 states) within the entire population but to accurately quantify diffusivity, the SpotOn web interface was used.

The Spot-On (26) analysis was performed on three frames or longer trajectories. The bound fraction and mean diffusion coefficient were extracted from the CDF of observed displacements over different time intervals. The cumulative displacement histograms were fitted with a 2-state model.

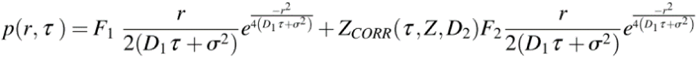

Where F_1_ and F_2_ are bound and free fractions, σ is single molecule localization error, D_1_ and D_2_ are diffusion coefficients of bound and free fractions, and Z_CORR_ is the correction factor for fast molecules moving out of the axial detection range (26). The axial detection range for JF646 on our setup is 294 nm. The following settings were used on the Spot-On web interface: bin width 0.01, number of time points 8, jumps to consider 4, use entire trajectories – No, Max jump (µm) 1. For model fitting, the following parameters were selected: D_bound_ (µm^2^/s) min 0.0001 max 0.5, D_free_ (µm^2^/s) min 0.5 max 5, F_bound_ min 0 max 1, Localization error (µm)-Fit from data-Yes min 0.01 max 0.1, dZ (µm) 0.65 for JF646, Use Z Correction-Yes, Model Fit CDF, Iterations 3.

### Indirect immunofluorescence

Cells were fixed with 4.5% formaldehyde for 2 hr at room temperature. The fixed cells were washed once with PBS and once with spheroplasting buffer (1.2 M sorbitol, 0.1 M phosphate buffer, pH 7.5) and resuspended in the same solution. Cells were spheroplasted using Zymolyase 20T in the presence of 25 mM β-Mercaptoethanol for 1hr at 30^°^ C. The spheroplasts were washed with a spheroplasting solution and transferred to poly-L-lysine-coated slides with Teflon-coated wells. The spheroplasts were flattened and permeabilized by immersing the slide in methanol for 5 min and acetone for 30 seconds. Blocking was done using 10 mg/ml BSA in PBS for 15 min. All the primary and secondary antibodies were diluted in the blocking reagent. Spheroplasts were incubated with primary antibodies for 1 h, washed thrice with PBS, and incubated with pre-adsorbed secondary antibodies for 1 h in the dark. Following further washes with PBS, samples were incubated with a mounting medium Vectashield (Victor laboratories, USA) for 15 min in the dark.

Antibody dilution: Rat anti-tubulin antibody (dilution: 1:200), TRITC conjugated Goat Anti Rat antibody (dilution: 1:200)

### Protein isolation and western blotting

Log phase culture was grown until OD_600_: 1. Cells were harvested and resuspended with 400 µl 0.1M NaOH for 30 minutes at room temperature. Cells were centrifuged, and the supernatant was discarded. Cell pellets were resuspended with 1X Laemmli sample buffer (2% SDS, 10% glycerol, 5% 2-mercaptoethanol, 0.002% bromophenol blue, and 60 mM Tris-HCl (pH -6.8)). An equal volume of 0.5 mm glass beads was added, the mixture was vigorously vortexed for 2 min, and incubated at 95 ^°^C for 10 minutes. Then, lysates were centrifuged in microfuge tubes at 13000 rpm for 10 minutes. Protein extracts were resolved on SDS– polyacrylamide gel, blotted onto PVDF membranes, and incubated with the following primary and secondary antibodies.

Antibody dilutions: Anti-HA Rabbit mAb (dilution: 1:2500), Anti-c-Myc (9B11) Mouse mAb (dilution: 1:2500), HRP conjugated Goat anti-rabbit antibody (dilution: 1:10000), HRP conjugated Goat anti-mouse antibody (dilution: 1:10000)

### Spot test

10-fold serial dilutions were made from the log phase cultures of all the strains at OD_600_ of 1. 2 µl of cell suspension from each dilution were placed on YPD Agar plates (and YPD agar + 1 mM auxin plates). Plates were incubated for 48 hr at 30 °C, and images were taken using the gel documentation system.

## RESULTS

### Strategy for SMIT of Ipl1 during different stages of mitosis

For visualizing single-molecules of Ipl1 in live cells of *S. cerevisiae*, we endogenously fused a protein self-labeling enzyme tag-HaloTag to the C-terminus of the *IPL1* gene for the controlled labeling of Ipl1. To identify the stages of the cell cycle and to mark the position of Ipl1 localization (kinetochores during metaphase, spindles during anaphase, Figure 1A), we expressed CloverGFP-Tub1. We used diploid strains for all the experiments with homozygous *pdr5*Δ, heterozygous *IPL1-HaloTag*, and heterozygous *CloverGFP-TUB1*. To check the functionality of the protein fusions and the overall viability of the genetically engineered strains, spot tests were performed with 10-fold serial dilution on YPD plates (Figure S1A). None of the strains showed significant growth delay compared to the parental strain. This result suggests that the protein fusions are functional and cells are healthy.

For sparse labeling of Ipl1-HaloTag, the log phase cells were treated with 30 nM JF646-HTL for 30 min and observed under the single-molecule imaging microscope (Figure S2A, Material and Methods). To quantify a broad range of kinetic behavior of Ipl1, time-lapse movies were acquired with two imaging regimes: ‘fast regime’ and ‘slow regime’ (Figure S2B, Material and Methods) (24).

Based on the CloverGFP-Tub1 morphology, the stage of mitosis (metaphase and anaphase) can be identified and the regions of interest (ROIs) can be defined for tracking the single molecules of Ipl1-HaloTag-JF646 at kinetochores and spindles (Figure 1C). Single-molecule tracking and data analysis were performed as described in Material and Methods (Figure S2) (24).

### Ipl1 is recruited to the kinetochores and spindles with different dynamics

For SMIT of Ipl1 at metaphase and anaphase, we used a diploid yeast strain with homozygous *pdr5*Δ, heterozygous *IPL1-HaloTag*, heterozygous *CloverGFP-TUB1* (YTK1804). The diploid strain was grown to the mid-log phase, the Ipl1-HaloTag was labeled with 30 nM of JF646-HTL for 30 min, and cells were imaged under the single-molecule imaging microscope using a slow imaging regime (Material and Methods, Figure S2A, S2B). The imaging and tracking were performed as described in Material and Methods.

We observed that the survival probability distributions for both stages fit well to the double exponential decay (Figure 2A, red line), suggesting two types of binding events: 1) binding with long residence time (slow fraction, shown by pink color in the pie chart) and 2) binding with short residence time (fast fraction, shown by blue color in the pie chart). The pink fraction represents specifically bound molecules of Ipl1 responsible for the phosphorylation of its substrates, whereas the blue fraction represents non-specifically/transiently bound molecules of Ipl1. Henceforth, we will discuss the specific bound fraction (pink fraction) only, as it represents Ipl1 molecules involved in substrate phosphorylation. In contrast, the blue and grey fractions do not contribute directly towards the Ipl1 function. To check that the pink fraction only represents a specific binding, we tracked Ipl1-HaloTag-JF646 in the cells where the CloverGFP-Tub1 signal is absent (at non-specific sites, Figure S3). The survival distribution fits well with the single-exponential decay, suggesting the absence of specific binding. We tracked Ipl1-HaloTag-JF646 at the kinetochores (during metaphase) and spindles (during anaphase, Figure 2A). Ipl1 showed residence times of 2.5±0.6 seconds and 9.5±0.5 seconds, respectively (pie charts, pink fractions, p<0.001). Also, the specific bound fraction (pink fraction) changes significantly between metaphase and anaphase (12.5±5.6% and 25.6±1.1%, respectively, p<0.001). These results suggest that Ipl1 is recruited to the kinetochores and spindles with different dynamics.

Several reports have shown three discrete binding sites (inner centromere, inner kinetochore, and outer kinetochore) of AK-B/Ipl1 during metaphase (11,12,25). While tracking Ipl1 at the kinetochores during metaphase, we could not differentiate the locations of these three sites due to the diffraction-limited resolution of the microscope. Hence, our metaphase tracking data represents the molecules at all the sites. To address this conundrum, we quantified the diffusion coefficients (D) of Ipl1 populations at the kinetochores during metaphase in cycling cells (Figure 2B) using the fast-imaging regime (Material and Methods) and Spot-On-based kinetic modelling (26). The presence of multiple populations of Ipl1 at the kinetochore should display multiple diffusion coefficients (as reported previously for the transcription pre-initiation complex components TBP and TFIIE) (17). However, we could not observe multiple populations of Ipl1 at the kinetochores. We used mean-squared displacement (MSD)-based logD histograms to identify the resolvable states (e.g., 2-states or 3-states) within the entire population (Figure 2B, left panel), which identified two populations only. However, this approach can produce unreliable quantification for mean D value because it depends on the choice of track length and linear fits to MSD. Therefore, the SpotOn-based kinetic modeling (26) was used to robustly quantify the mean D value. This analysis showed a 48% bound fraction with a mean D value of 0.08 µm^2^/s (Figure 2B, right panel). This result suggests that despite having multiple discrete sites for the localization of Ipl1 at the kinetochores, all the molecules of Ipl1 at the kinetochore have the same diffusion characteristics. To support this result, two recent studies have demonstrated that the inner centromere and the inner kinetochore CPC targeting mechanisms are at least partially redundant for chromosome biorientation and cell viability in budding yeast (8,9). Additionally, mutations in the SAH domain of Sli15 prevent CPC localization to all three sites in budding yeast (12). All this evidence suggests that despite having three discrete sites for Ipl1 localization at the kinetochores, all these sites may be functionally redundant.

### The absence of Ctf19 or Bub1 reduces the specific bound fraction of Ipl1 at the kinetochores and increases the target search time

As mentioned in the introduction, Ipl1 is recruited to the kinetochore as a part of the CPC by two pathways: A) Bub1/Sgo1 mediated recruitment, and B) Ctf19 (COMA complex) mediated recruitment (Figure 1B). Both these mechanisms work redundantly to promote chromosome biorientation. Hence, we want to address how the absence of any of these mechanisms affects the recruitment dynamics of Ipl1 at the kinetochore. We created homozygous deletions of *BUB1* or *CTF19* in the diploid strain used above (GMY009, GMY303). The cells were labeled and imaged for SMIT, and the data analysis was performed as described before (Figure 2A).

We observed that the absence of Ctf19 reduces the specific bound fraction of Ipl1 at the kinetochore significantly (from 12.5±5.6% (Figure 2A) to 5.3±1.7%, p<0.05 (Figure 2C)), without altering its residence time significantly (from 2.5±0.6 seconds (Figure 2A) to 3.6±0.5 seconds (Figure 2D), p=not significant). As Ctf19 is the known recruiter of Ipl1 to the kinetochores, we asked if Ipl1 takes a long time to find the kinetochore in its absence. We calculated the search time (τ_search_, the time required by Ipl1 to reach the kinetochores, Figure S2E, Material and Methods). The absence of Ctf19 increases the search time of Ip1l to find the kinetochores, from 9.6±5.1 seconds (Figure 2A metaphase) to 20.6±4.2 seconds (Figure 2C, *ctf19*Δ, p<0.001). Conversely, the absence of Bub1 completely abolished the specific bound fraction of Ipl1 at the kinetochore (Figure 2C), suggesting the dominance of the Bub1-mediated recruitment of Ipl1 at the kinetochores over the Ctf19-mediated recruitment. Both these pathways are not entirely independent. Without Bub1, the Ctf19-mediated pathway cannot recruit Ipl1 to the kinetochores.

### Tension across the kinetochores diminishes Ipl1 binding to the kinetochores

Ipl1 is best known for its role in tension sensing at the kinetochores during metaphase for error correction (27–29). So, we asked how the tension across the kinetochore alters the dynamic recruitment of Ipl1 during metaphase. We designed an experimental strategy by which we can keep all the kinetochores under tension, followed by adding a microtubule depolymerizing drug (benomyl) to release the tension. We used an auxin-inducible degron system (Material and Methods) for the conditional depletion of the activator of the anaphase-promoting complex, Cdc20. By depleting Cdc20, cells were arrested at metaphase with all the kinetochores attached to the spindles, and they were under tension due to the pulling forces exerted by the spindles (Figure 3A). For this purpose, we genetically engineered a diploid yeast strain with homozygous *pdr5*Δ, homozygous *CDC20-AID*-6HA,* heterozygous *IPL1-HaloTag*, heterozygous *NDC10-GFP*, and heterozygous *pADH-AFB2* (YTK1807). For this experiment, we used *Ndc10-GFP* instead of CloverGFP-Tub1 because microtubule depolymerization may lead to the disappearance of CloverGFP-Tub1, which may increase inaccuracies in ROI generation for tracking. Ndc10 is an inner kinetochore protein that directly visualizes the position of the metaphase kinetochores in the presence and absence of microtubules. It also facilitates ROI generation for tracking. SMIT was performed for Ipl1-HaloTag-JF646 after adding auxin for 2 hr to the log phase cells (Figure S2A, S2B). We added benomyl at 90 µg/ml concentration to release the tension and incubated the cells for an additional 1 hr (Figure S2A). We performed immunofluorescence analysis using anti-tubulin antibodies and DAPI from both these samples to confirm the metaphase arrest upon depleting Cdc20 and microtubule depolymerization upon benomyl treatment. Immunofluorescence analysis confirmed that >85% of cells showed metaphase spindle upon depleting Cdc20 and the absence of elongated microtubules in >95% of cells after treatment with benomyl (Figure S4).

Our SMIT data demonstrated that the dynamic recruitment of Ip1l diminished at the kinetochores when the kinetochores were under tension (Figure 3B), as seen by the absence of the specifically bound fraction of Ipl1. Upon releasing the tension, Ipl1 re-localized to the kinetochores (19.9±3.4% pink fraction with a residence time of 3±0.4 seconds, Figure 3B). This result aligns with a recent report in which authors have demonstrated the same phenomenon using Ipl1-3xGFP (27). This data suggests that the tension across the kinetochores can modulate the dynamic recruitment of Ipl1 at the kinetochore.

### Conditional depletion of Glc7 increases the molecular crowding of Ipl1 at the metaphase kinetochores

Glc7 (PP1) antagonizes Ipl1-mediated phosphorylation (14). Ipl1 and Glc7 modulate the phosphorylation of a common set of substrates (Ndc10, Dam1, H3). However, the precise interplay between the kinase and phosphatase needs to be better understood. To address this question, we quantified the single-molecule dynamics of Ipl1 at the kinetochore in the absence of Glc7 during metaphase. We genetically engineered a diploid yeast strain with homozygous *pdr5*Δ, homozygous *Glc7-AID*-6HA,* heterozygous *IPL1-HaloTag*, heterozygous *CloverGFP-TUB1,* and heterozygous *pADH-AFB2* (GMY043). SMIT was performed for Ipl1-HaloTag-JF646 with and without auxin treatment for 2 hr (Figure S2A).

We observed that in the absence of Glc7, the fraction of specific bound molecules of Ipl1-HaloTag-JF646 increases from 12.5±5.6% to 19.6±4.3% (p<0.001, Figure 3C) and the residence time increases from 2.5±0.6 seconds to 5.3±1 seconds, (p<0.005, Figure 3C). This result suggests that the absence of Glc7 increases the retention of Ipl1 at the kinetochores. Hence, the presence of Glc7 is required for the fast turnover/exchange of the Ipl1 at the kinetochores during metaphase. Rapid turnover of Ipl1 at the metaphase kinetochores may keep Glc7 away from their common substrates to keep them phosphorylated.

### SMIT of CPC components reveals the hierarchical assembly of the CPC at the kinetochores during metaphase

Ipl1 is recruited to the kinetochores during metaphase as a part of the CPC (30). It is unknown how these four-membered protein complexes assemble at the kinetochores. To address this question, we designed an experiment to track the single-molecule dynamics of other members of the CPC (Bir1, Nbl1, Sli15). The hypothesis is that if the entire complex is recruited to the kinetochores dynamically, all components’ residence time will be comparable. Otherwise, if there is a hierarchical assembly, all these components will show different residence times, depending on their order in the assembly process. The component recruited first will show the longest residence time, compared to the component recruited last. For this experiment, we engineered four different strains with heterozygous CloverGFP-Tub1, homozygous *pdr5*Δ, and heterozygous C-terminal-HaloTag fusions to the CPC components (Bir1, Nbl1, Sli15, and Ipl1; GMY013, GMY014, GMY044, and GMY 1804, respectively). We confirmed the functionality of the C-terminal fusions by the spot test (Figure S1A) and could not observe significant growth defects.

SMIT was performed for all four diploid strains, as mentioned above. We observed that the residence times of Bir1 and Nbl1 are comparable (6.4±0.6 seconds and 5.5±0.6 seconds, respectively, p=not significant), and that of Sli15 and Ipl1 are comparable (2.1±0.7 s and 2.5±0.6 s, respectively, p=not significant). This result suggests that Bir1-Nbl1 and Sli15-Ipl1 are recruited to the kinetochores as complexes. However, there is a significant difference in the residence time of the Bir1-Nbl1 complex and the Sli15-Ipl1 complex (p<0.001). This result suggests the hierarchical assembly of the CPC components to the kinetochores during metaphase, Bir1-Nbl1 heterodimer is recruited first to the kinetochore, followed by Sli15-Ipl1 heterodimer.

**Figure 4:**
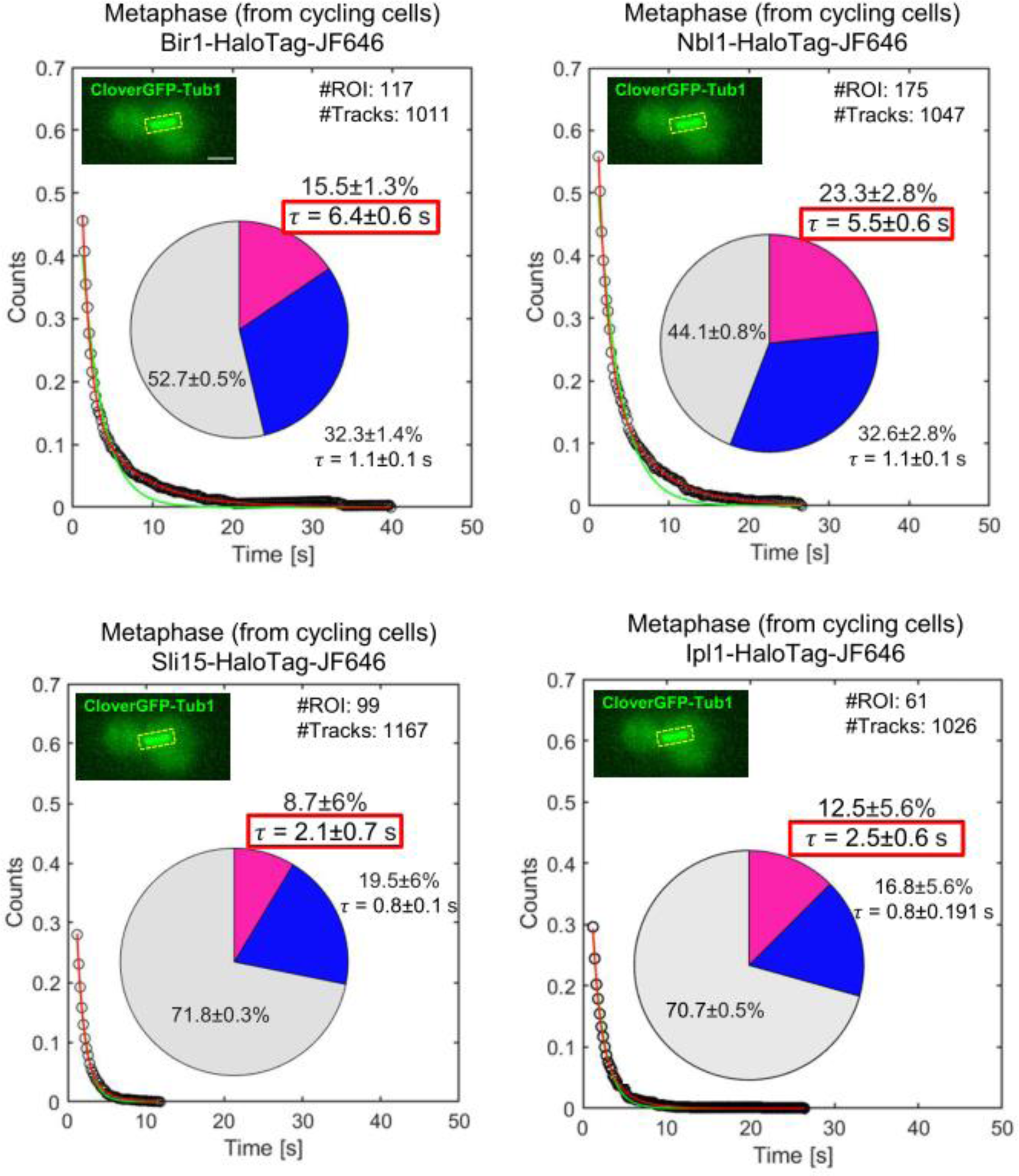
Hierarchical assembly of the CPC at the kinetochore revealed by SMIT of Bir1, Nbl1, Sli15, and Ipl1 at metaphase: SMIT of Bir1-HaloTag-JF646, NBl1-HaloTag-JF646, Sli15-HaloTag-JF646, and Ipl1-HaloTag-JF646 during metaphase using a slow imaging regime. Survival probability distributions and pie charts are represented as Figure 2A. “#Tracks” represents the total number of tracks analyzed, and “#ROIs” represents the total number of cells tracked. Scale: 2 µm. The inset images show the ROIs used for tracking. The data shown for Ipl1-HaloTag-JF646 is the same as it is shown in Figure 2A (for metaphase). However, it has been shown again for easy comparison with the other CPC subunits.

## DISCUSSION

AK-B in humans and Ipl1 in yeast *S. cerevisiae* have been studied for a few decades as they are the most promising targets for cancer therapies (31). Throughout these years, several ensemble averaging methods such as ChIP, immunofluorescence, live cell imaging, western blotting, and proteomic analysis have been employed to understand the dynamics and functions of AK-B/Ipl1. However, all these methods provided a static picture of AK-B/Ipl1 localization and function due to population-based averaging. The dynamic information was lost. SMIT has emerged as a powerful technique to visualize and quantify the dynamics of proteins in live cells with high spatiotemporal resolution (18). Here, we employed this method to quantify the recruitment dynamics of Ipl1 in yeast *S. cerevisiae* and to understand how it changes in the absence of several modulators.

Using this method, we could quantify the time-scale of Ipl1 activities, such as how long it takes to phosphorylate its substrates (e.g. 2.5±0.6 seconds at metaphase, 9.5±0.5 seconds at anaphase), how long it takes to find its target sites (kinetochores) during metaphase (e.g. 9.6±5.1 seconds in wild-type, 20.6±4.2 seconds in the absence of Ctf19), and what fraction of molecules are involved in phosphorylation (e.g. 12.5±5.6% during metaphase, 25.6±1.1% during anaphase). Intriguingly, Ipl1 spends less time at the kinetochores (2.5±0.6 seconds) compared to spindles (9.5±0.5 seconds), though it has more substrates to phosphorylate at the kinetochores (Cse4, Ndc10, Ndc80/Hec1, Dam1, Dsn1) compared to spindles (Cin8, Bim1, Ase1). Hence, the rapid turnover at the kinetochore may be necessary for its kinetochore functions (kinetochore assembly and spindle checkpoint). It may create a molecular crowd around kinetochores that keeps the phosphatase (Glc7) away so the substrates of Ipl1 at the kinetochore remain phosphorylated. A recent report from Tanaka lab shows that rapid turnover of Ipl1 at the kinetochore is not required to achieve bi-orientation (32). These experiments were done in metaphase-arrested cells (by depleting Cdc20) and using artificial tethering of Ipl1-Sli15 to the kinetochore proteins Mif2 and Ndc80. So, it is possible that in cycling cells, rapid turnover of Ipl1 at the kinetochore is essential to keep the Glc7 away to support other functions such as kinetochore assembly and spindle checkpoint. Our data also support this hypothesis, as in the absence of Glc7, the residence time and the specific bound fraction of Ipl1 increases at the kinetochores during metaphase (from 2.5±0.6 seconds to 5.3±1 seconds and from 12.5±5.6% to 19.6±4.3%, Figure 3C). We have also shown that the tension across the kinetochores evicts Ipl1 from the kinetochore, consistent with the recent report (27). The absence of Ctf19 reduces the specific bound fraction of Ipl1 without changing the residence time significantly. This result suggests that Ctf19 is not essential for the rapid turnover of the Ipl1 at the kinetochores. However, it provides a platform for the specific binding of Ipl1 to the kinetochore. In the absence of Bub1, the specific binding of Ipl1 to the kinetochores was abolished, suggesting the absence of the Ctf19-mediated recruitment of Ipl1 to the kinetochore. Hence, neither of these pathways of Ipl1 recruitment to the kinetochores is mutually exclusive. SMIT of CPC components showed hierarchical assembly of the CPC at the kinetochore, Bir1-Nbl1 being the first to be recruited at the kinetochores, followed by the Sli15-Ipl1 complex.

The method we have developed here for the SMIT of Ipl1 in *S. cerevisiae* can be used to understand the dynamics of several other mitotic kinases, phosphatases, or any other proteins in yeast and other model systems. It is in infancy to understand the dynamic interplay between a kinase and a phosphatase for maintaining a critical phosphorylation level of their substrates. So, this method can be a valuable tool to understand such dynamic biological processes. Recent reports have demonstrated the liquid-liquid phase separation (LLPS) phenomenon for the CPC components Survivin, Borealin, and INCENP *in vitro* (33–34). Also, it has been demonstrated by live-cell single-molecule imaging that the LLPS accelerates the target-search process for transcription machinery (35). Hence, it will be interesting to address in the future if the CPC components show LLPS behavior in live yeast. The SMIT-based assay may provide evidence for the LLPS behavior of CPC components in live cells. Understanding the dynamic regulation of AK-B/Ipl1 may open new avenues for drug development by which its recruitment dynamics can be modulated instead of a widely used approach of inhibiting its kinase activity.

This study has two limitations. 1) Ipl1 phosphorylates several proteins at the kinetochores during metaphase (Cse4, Ndc10, Dam1, Ndc80/Hec1, Dsn1). We cannot distinguish the binding dynamics of Ipl1 for phosphorylating each of these proteins. So, the residence time and the other parameters quantified for Ipl1 dynamics at the kinetochores are averaged out for all its kinetochore substrates. 2) Due to the diffraction-limited resolution of the microscope, we cannot distinguish the three discrete sites of Ipl1 localization at the metaphase kinetochores. Previous reports have shown three discrete sites using immunofluorescence in cell lines (11). So, the estimation of residence time at the kinetochore represents the residence times of all the molecules of Ipl1 localized at three distinct sites.

## DATA AVAILABILITY

The data underlying this article are available in Mendeley Data, at https://data.mendeley.com/preview/hbfm65dg6v?a=ec827291-ccbb-4cf8-9848-ed784546cca4

## AUTHOR CONTRIBUTIONS

Nitesh Kumar Podh: Methodology, Software, Validation, Formal Analysis, Investigation, Data Curation, Writing-Original Draft, Writing-Review and Editing, Visualization Ayan Das: Methodology, Software, Validation, Formal Analysis, Investigation, Data Curation, Writing-Original Draft, Writing-Review and Editing, Visualization Aakriti Kumari: Methodology, Software, Investigation, Formal Analysis, Writing-Review and Editing Kirti Garg: Methodology, Software, Investigation, Formal Analysis, Writing-Review and Editing Rashmi Singh Yadav: Methodology, Software, Investigation, Formal Analysis, Writing-Review and Editing Kirti Kashyap: Software, Validation, Formal Analysis, Writing-Review and Editing Sahil Islam: Software, Validation, Formal Analysis, Writing-Review and Editing Anupam Gupta: Software, Validation, Formal Analysis, Writing-Review and Editing, Visualization, Supervision, Project Administration Gunjan Mehta: Conceptualization, Resources, Writing-Original Draft, Writing-Review and Editing, Visualization, Supervision, Project Administration, Funding Acquisition

## ACKNOWLEDGEMENTS

We are grateful to Dr. David Ball (National Cancer Institute (NCI), National Institutes of Health (NIH), USA) for providing support with the data analysis.

