## Supplementary Information for "Single-molecule tracking reveals the dynamics of Ipl1 recruitment to the kinetochores and spindles in *S. cerevisiae*"

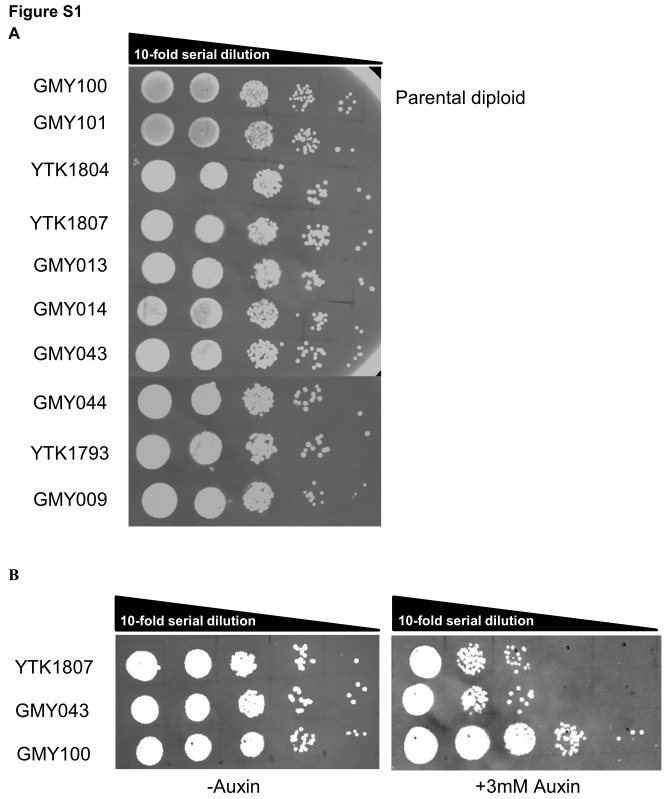
**Supplementary Information**

**Figure S1** **Spot tests to check the functionality of all the C-terminal fusions and viability of yeast cells: (A)** 10-fold serial dilutions of all the diploid strains used in this study were plated on YPD plates. GMY100 was used as a parental strain (without any genetic engineering). **(B)** Functional assay for auxin-inducible depletion of the Cdc20-AID* and Glc7-AID*. 10-fold serial dilutions were plated on YPD and YPD+3 mM Auxin plates. GMY100 was used as a parental strain (without any genetic engineering).

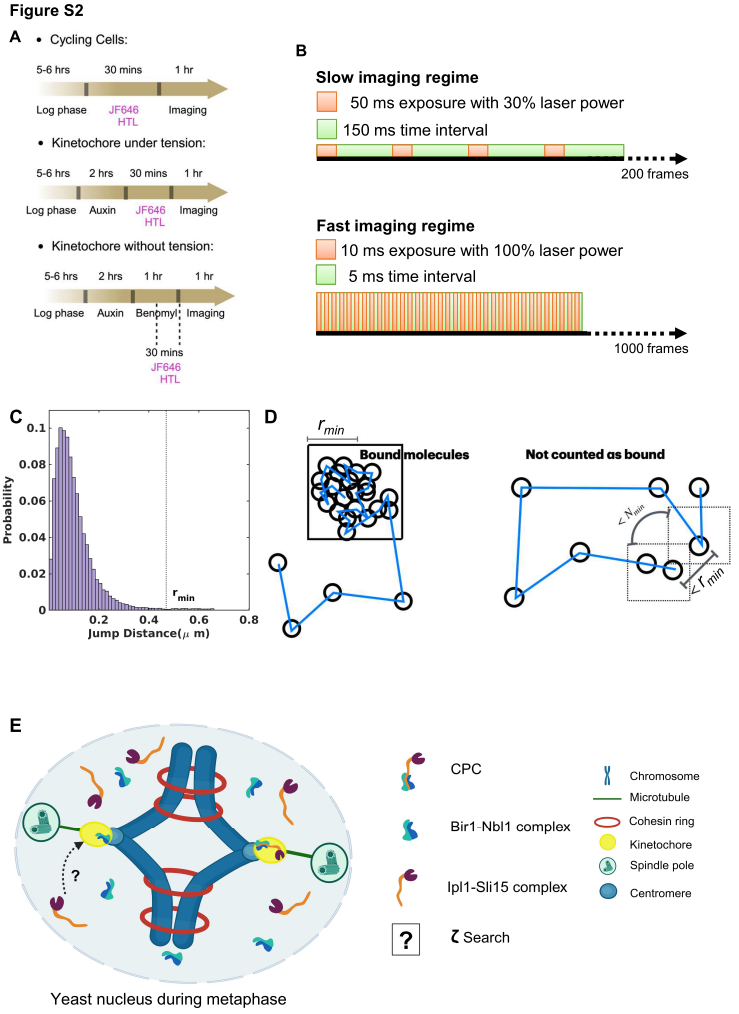

**Figure S2 Culturing, imaging, and tracking strategies:** **(A)** Culturing regime: Overnight grown cells were inoculated in a fresh CSM media and grown for 5 hr (for log phase). Cells were treated with JF646-HTL for 30 min, washed twice with CSM, and imaged under the microscope. When necessary, auxin treatment was given 2 hr before the addition of JF646-HTL. When necessary, benomyl treatment was given for 1 hr after auxin treatment, along with incubation with JF646-HTL for 30 min. In all cases, the cells were imaged for 1 hr from the same agarose pad. Every 1 hr, fresh samples were taken. **(B)** Imaging regimes: Time-lapse movies were acquired with a ‘slow imaging regime’ (200 ms time interval for 200 frames, 50 ms exposure time) to estimate dwell time, whereas, time-lapse movies were acquired with a ‘fast imaging regime’ (15 ms interval for 1000 frames, 10 ms exposure time) to estimate diffusion parameters (fraction of bound and unbound molecules, and diffusion coefficients). **(C)** SMIT was performed for chromatin-bound histone H3 (Hht1-HaloTag-JF646) to quantify the value of *r_min_*. Frame-to-frame displacements were plotted as a histogram. The *r_min_* value defines the maximum frame-to-frame distance traveled by 99% of histone H3 molecules. **(D)** Schematics to explain the terms: *r_min_* and *N_min._ r_min_* is the maximum frame-to-frame distance traveled by 99% histone H3 molecules in time-lapse movies acquired with a 200 ms time interval. *N_min_* is the minimum number of frames for which a bound molecule should not travel more than *r_min_* distance frame-to-frame. **(E)** Schematics of the target-search mechanism of CPC to find kinetochores during metaphase. τ_search_ is the time required by Ipl1 to reach the kinetochore. As demonstrated in Figure 4, the assembly of CPC at the kinetochores takes place in the hierarchical manner, the Bir1-Nbl1 complex assembles first, followed by the recruitment of the Sli15-Ipl1 complex.

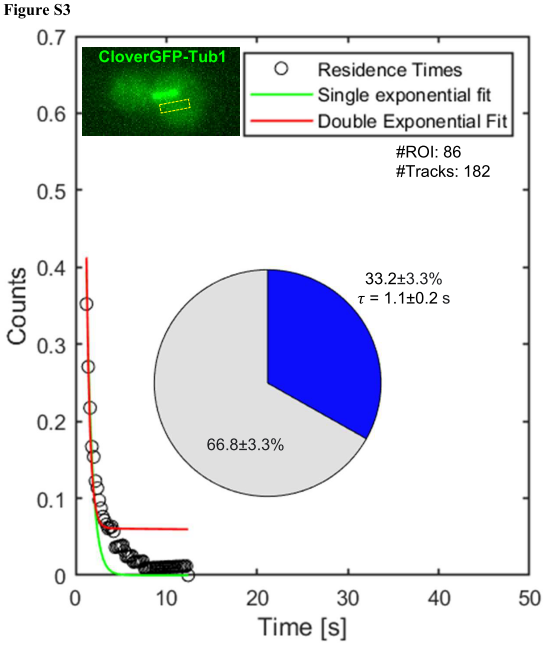

**Figure S3:** **Tracking of Ipl1-HaloTag-JF646 at the non-specific site in the yeast cells:** From the same set of images used for tracking Ipl1-HaloTag-JF646 at metaphase (Figure 2A), molecules were tracked by making ROIs away from the SPBs, kinetochores and spindles (at non-specific sites). Survival probability distribution fit well with the single exponential decay curve (green curve). Hence, the pie chart represents only the fraction of molecules bound with short dwell time (green fraction), and diffusing molecules (grey fraction). “n” represents the total number of tracks analyzed, and “#ROIs” represents the total number of cells tracked. The inset images show the ROI (red) used for tracking. Cell boundaries are drawn manually. Scale: 2 µm

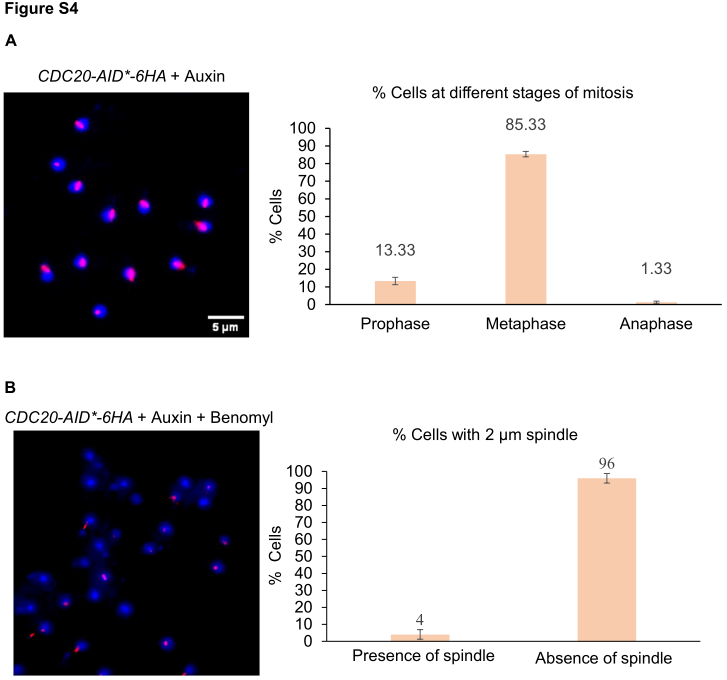

**Figure S4: Immunofluorescence to validate metaphase arrest and microtubule depolymerization:** **(A)** Immunofluorescence was performed using anti-tubulin antibodies and DAPI staining to determine the stages of the cell cycle. After 2.5 hr of auxin treatment (for Cdc20 depletion), >85% of cells showed 2 µm long tubulin staining indicating the metaphase stage. **(B)** Upon microtubule depolymerization (by adding benomyl for 1 hr), loss of 2 µm long tubulin structures was observed in >95% of cells, confirming microtubule depolymerization. The single foci in some of the nuclei represent SPBs, where the monomers of tubulin are present.

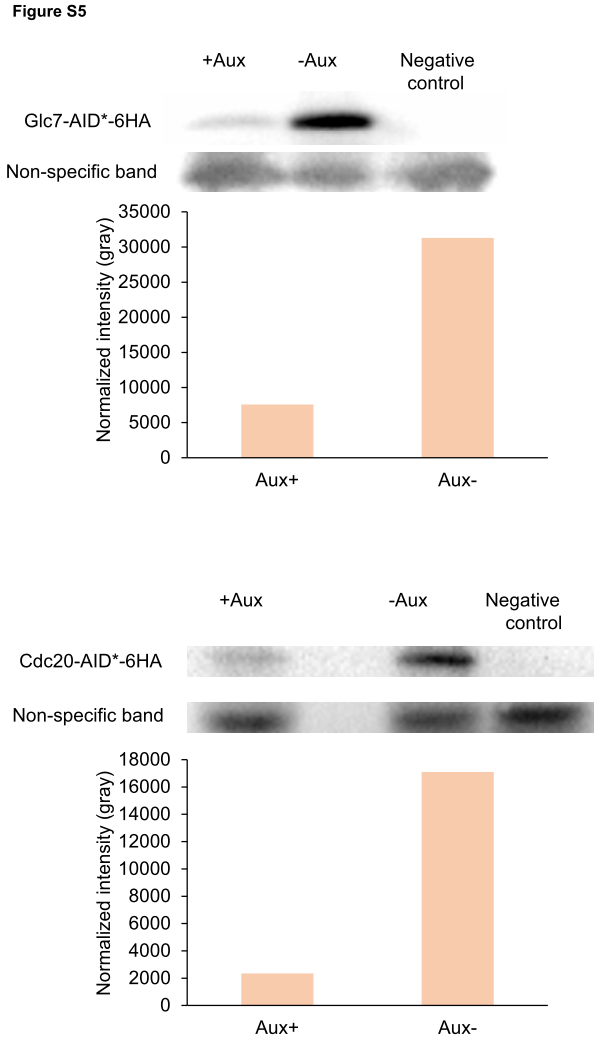

**Figure S5:** **Western blotting to confirm the auxin-induced depletion of Glc7 and Cdc20:** The addition of auxin depleted Glc7 and Cdc20 significantly (>4 times for Glc7 and >8 times for Cdc20). A non-specific band on the western blot was used as a loading control.

**Movie S1:** Single-molecule imaging of Ipl1-HaloTag-JF646 at metaphase and anaphase with 200 ms time interval, 50 ms exposure, and 30% laser power. CloverGFP-Tub1 was used as a localization marker of Ipl1 and to identify the stages of mitosis. Left: the single-focal plane image of CloverGFP-Tub1, middle: the single-focal plane time-lapse movie of Ipl1-HaloTag-JF646, right: Merged channels. Playback speed: real speed (5 fps)

**Movie S2:** Single-molecule imaging of Ipl1-HaloTag-JF646 at metaphase kinetochores with 200 ms time interval, 50 ms exposure, and 30% laser power. Here the kinetochores are under tension due to conditional depletion of Cdc20. So, the single-molecules of Ipl1-HaloTag-JF646 are visible, diffusing near the kinetochores but they do not show binding. CloverGFP-Tub1 was used as a localization marker of Ipl1 and to identify the stages of mitosis. Left: the single-focal plane image of CloverGFP-Tub1, middle: the single-focal plane time-lapse movie of Ipl1-HaloTag-JF646, right: Merged channels. Playback speed: real speed (5 fps)

**Table S1. List of diploid yeast strains used for this study:**

All the yeast strains are derived from BY4741 (MATa) and BY4742 (MATalpha) (S288C background, Research Genetics/Invitrogen, USA).

| **Sr. No** | **Strain ID** | **Genotype** | **Source** |
| --- | --- | --- | --- |
| 1. | GMY009 | *his3-d1 leu2-d0 met15-d0 ura3-d0 pdr5Δ::LoxP trp1d::pADH-AFB2-LEU2 CDC20-AID*-9Myc-HIS3 IPL1-HaloTag-TRP1 ctf19Δ::hphNT1*  *his3-d1 leu2-d0 lys2-d0 ura3-d0 pdr5Δ::LoxP CDC20-AID*-9Myc-HIS3 tub1Δ::pHIS3p:CloverGFP-TUB1+3'UTR-URA3 ctf19Δ::hphNT1* | This study |
| 2. | GMY013 | *his3-d1 leu2-d0 met15-d0 ura3-d0 pdr5Δ::LoxP trp1d::pADH-AFB2-LEU2 CDC20-AID*-9Myc-HIS3 BIR1-HaloTag-TRP1*  *his3-d1 leu2-d0 lys2-d0 ura3-d0 pdr5Δ::LoxP CDC20-AID*-9Myc-HIS3 tub1Δ::pHIS3p:CloverGFP-TUB1+3'UTR-URA3* | This study |
| 3. | GMY014 | *his3-d1 leu2-d0 met15-d0 ura3-d0 pdr5Δ::LoxP trp1d::pADH-AFB2-LEU2 CDC20-AID*-9Myc-HIS3 NBL1-HaloTag-TRP1*  *his3-d1 leu2-d0 lys2-d0 ura3-d0 pdr5Δ::LoxP CDC20-AID*-9Myc-HIS3 tub1d::pHIS3p:CloverGFP-TUB1+3'UTR-URA3* | This study |
| 4. | GMY043 | *his3-d1 leu2-d0 met15-d0 ura3-d0 pdr5Δ::LoxP trp1d::pADH-AFB2-LEU2 CDC20-AID*-9Myc-HIS3 IPL1-HaloTag-TRP1 GLC7-AID*-6HA-hphNT1*  *his3-d1 leu2-d0 lys2-d0 ura3-d0 pdr5Δ::LoxP CDC20-AID*-9Myc-HIS3 tub1d::pHIS3p:CloverGFP-TUB1+3'UTR-URA3 GLC7-AID*-6HA-hphNT1* | This study |
| 5. | GMY044 | *his3-d1 leu2-d0 met15-d0 ura3-d0 pdr5Δ::LoxP trp1d::pADH-AFB2-LEU2 CDC20-AID*-9Myc-HIS3 SLI15-HaloTag-TRP1*  *his3-d1 leu2-d0 lys2-d0 ura3-d0 pdr5Δ::LoxP CDC20-AID*-9Myc-HIS3 tub1d::pHIS3p:CloverGFP-TUB1+3'UTR-URA3* | This study |
| 6. | GMY100 | *his3-d1 leu2-d0 met15-d0 ura3-d0 his3-d1 leu2-d0 lys2-d0 ura3-d0* | This study |
| 7. | YTK1793 | *his3-d1 leu2-d0 met15-d0 ura3-d0 HHT1-HaloTag-URA3 pdr5Δ::LoxP-hphNT1-LoxP*  *his3-d1 leu2-d0 lys2-d0 ura3-d0 ace1Δ::URA3 Cu1::(LacO)256 TRP1::pHis3-GFP-FFAT-LacI-NLS- HIS3 pdr5Δ::LoxP-LEU2-LoxP* | This study |
| 8. | YTK1804 | *his3-d1 leu2-d0 lys2-d0 ura3-d0 pdr5Δ::LoxP CDC20-AID*-9Myc-HIS3 tub1d::pHIS3p-CloverGFP-TUB1+3'UTR-URA3*  *his3-d1 leu2-d0 met15-d0 ura3-d0 pdr5Δ::LoxP trp1d::pADH-AFB2-LEU2 CDC20-AID*-9Myc-HIS3 IPL1-HaloTag-TRP1* | This study |
| 9. | YTK1807 | *his3-d1 leu2-d0 met15-d0 ura3-d0 pdr5Δ::LoxP trp1d::pADH-AFB2-LEU2 CDC20-AID*-9Myc-HIS3 IPL1-HaloTag-TRP1*  *his3-d1 leu2-d0 lys2-d0 ura3-d0 pdr5Δ::LoxP CDC20-AID*-9Myc-HIS3 NDC10-3xGFP-URA3* | This study |
| 10. | GMY303 | *his3-d1 leu2-d0 lys2-d0 ura3-d0 pdr5-d::LoxP CDC20-AID*-9Myc::HIS3 tub1d::pHIS3p:CloverGFP-TUB1+3'UTR::URA bub1-d::KANMX*  *his3-d1 leu2-d0 met15-d0 ura3-d0 pdr5-d::LoxP trp1d::pADH-AFB2::LEU2 CDC20-AID*-9Myc::HIS3 IPL1-HaloTag::TRP1 bub1-d::NATMX* | This study |

**Table S2: List of plasmids used in this study:**

| **Sr No.** | **Plasmid name** | **Description** | **Source** |
| --- | --- | --- | --- |
| 1 | pTSK573 | For C-terminal fusion of the *HaloTag* using *TRP1* selection marker | [Addgene: 190881](https://www.addgene.org/190881/) |
| 2 | *pHyg-AID*-6HA* | For C-terminal fusion of the *AID* (Auxin-Inducible Degron) tag for the conditional depletion of a protein of interest gene, selection marker: Hygromycin B | [Addgene: 99520](https://www.addgene.org/99520/) |
| 3 | pAG32 | For gene deletion using hygromycin B selection marker | [Euroscarf: P30106](http://www.euroscarf.de/plasmid_details.php?accno=P30106) |
| 4 | *pHIS3p:CloverGFP-TUB1+3’UTR::URA3* | For the expression of CloverGFP-Tub1 as a localization marker | [Addgene: 50636](https://www.addgene.org/50636/) |
| 5 | pTSK559 | For the expression of *AFB2* from *ADH1* promoter | (20) |
| 6 | pTSK405 | For C-terminal fusion with *3xGFP* tag (*NDC10-3xGFP*, *IPL1-3xGFP*) | (20) |

**Table S3: List of primers used in this study:**

Homologous sequences for site-specific integration/recombination are shown in blue fonts. Sequences homologous to plasmid DNA and genomic DNA are shown in red and black fonts, respectively.

| **Sr No.** | **Primer Id** | **Description** | **Template** | **Sequence** |
| --- | --- | --- | --- | --- |
| 1 | GM29 | Forward primer for *SLI15-HaloTag-TRP1* | pTSK573 | *ATCCTAGGCTAAACAGGTTGAAACCGCGTCAAATTGTGCCCAAAAGGTCTCAAGCGGCCGCCGCTGCTGC* |
| 2 | GM30 | Reverse primer for *SLI15-HaloTag-TRP1* | pTSK573 | *ATTTAATGTTAACCAGTTTGAATTTTTCTTTTCTGGGGTAATCGAATTCACTATTTCTTAGCATTTTTGACG* |
| 3 | GM31 | Diagnostic forward primer for *SLI15-HaloTag-TRP1* | *S. cerevisiae* genomic DNA | *GGCTAGCGTAACTTTAGCGG* |
| 4 | GM32 | Diagnostic reverse primer for *SLI15-HaloTag-TRP1* | *S. cerevisiae* genomic DNA | *CGTAATGCAGGGGGAATACC* |
| 5 | GM33 | Forward primer for *BIR1-HaloTag-TRP1* | pTSK573 | *TTTGGAAGATGACAATCAATTGATCGATATTGCTAAGAAAATGGGCATTTTACAAGCGGCCGCCGCTGCTGC* |
| 6 | GM34 | Reverse primer for *BIR1-HaloTag-TRP1* | pTSK573 | *AAACTACAAAAAATACAAACCTTTAGCCTGTTTATCAAATTAGTTAGCTACTATTTCTTAGCATTTTTGACG* |
| 7 | GM35 | Diagnostic forward primer for *BIR1-HaloTag-TRP1* | *S. cerevisiae* genomic DNA | *GCAGAAGAGTTGGACATGAC* |
| 8 | GM36 | Diagnostic reverse primer for *BIR1-HaloTag-TRP1* | *S. cerevisiae* DNA | *CAGATATCTGCGATGCGGCG* |
| 9 | GM37 | Forward primer for *NBL1-HaloTag-TRP1* | pTSK573 | *CAAAGGAACTAATCAGAGAGGTGCTTGAACAGGAAGGACGCCGTATAGAACAAGCGGCCGCCGCTGCTGC* |
| 10 | GM38 | Reverse primer for *NBL1-HaloTag-TRP1* | pTSK573 | *AGGTGCATCATTGCGAATACCGAGAAAGGGTCCATTATACGAACTAATCACTATTTCTTAGCATTTTTGACG* |
| 11 | GM39 | Diagnostic forward primer for *NBL1-HaloTag-TRP1* | *S. cerevisiae* genomic DNA | *CTCTACCCCCAACCTTCACC* |
| 12 | GM40 | Diagnostic reverse primer for *NBL1-HaloTag-TRP1* | *S. cerevisiae* genomic DNA | *GGGATGTAACCACACGCTGC* |
| 13 | GM41 | Forward primer for *GLC7-AID*-6HA-hphNT1* | *pHyg-AID*-6HA* | *AGCCAGCCCAAAAAAGTCTACCAAGGCAAGCTGGGGGTAGAAAGAAAAAACGTACGCTGCAGGTCGAC* |
| 14 | GM42 | Reverse primer for *GLC7-AID*-6HA-hphNT1* | *pHyg-AID*-6HA* | *TAATAAGTATTTTCCTTTTTAAACTTTGATTTAGGACGTGAATCTATTTAATCGATGAATTCGAGCTCG* |
| 15 | GM43 | Diagnostic forward primer for *GLC7-AID*-6HA-hphNT1* | *S. cerevisiae* genomic DNA | *CGCTGGTGCAATGATGAGTG* |
| 16 | GM79 | Diagnostic reverse primer for *GLC7-AID*-6HA-hphNT1* | *S. cerevisiae* genomic DNA | *GACGAGTGATGATTGCATCTTCC* |
| 17 | GM49 | Forward primer for *ctf19*Δ | pAG32 | *CTTGGAGCTAGTGTGATCTTGTTGATACTAGGTCGGCAAAGAACGCAAATGCCAGCTGAAGCTTCGTACG* |
| 18 | GM50 | Reverse primer for *ctf19*Δ | pAG32 | *TATCGGAATCGTTTAAGCAAGCCGTCCAGTTGGCAATGGCAAATGGAACAGGCCACTAGTGGATCTG* |
| 19 | GM51 | Diagnostic forward primer for *ctf19*Δ | *S. cerevisiae* genomic DNA | *GGTAATGTAACCGGTAATGG* |
| 20 | GM52 | Diagnostic reverse primer for *ctf19*Δ | *S. cerevisiae* genomic DNA | *CGACGATGCAAATGAATTGC* |
| 21 | GM66 | Forward primer for *pdr5*Δ | pAG32 | *AGACCCTTTTAAGTTTTCGTATCCGCTCGTTCGAAAGACTTTAGACAAAAGCCAGCTGAAGCTTCGTACG* |
| 22 | GM67 | Reverse primer for *pdr5*Δ | pAG32 | *AAATTCAAGAAAATTGAAATGTAGAAAGCTCGCTGAATTAAGAAAAAAAAGGCCACTAGTGGATCTG* |
| 23 | GM68 | Diagnostic forward primer for *pdr5*Δ | *S. cerevisiae* genomic DNA | *CTCTTCTACGCCGTGGTACG* |
| 24 | GM69 | Diagnostic reverse primer for *pdr5*Δ | *S. cerevisiae* genomic DNA | *GAAGACGGTTCGCCATTCG* |
| 25 | GM70 | Forward primer for *IPL1-HaloTag-TRP1* | pTSK573 | *TGCATCCTTGGATACTAAGAAACATGCCCTTTTGGGAAAATAAGCGGTTACAAGCGGCCGCCGCTGCTGC* |
| 26 | GM71 | Reverse primer for *IPL1-HaloTag-TRP1* | pTSK573 | *GGGAGTGATTAATAGTGCCCTTCAAACGATTCTGTCATACTTTAATTCTACTATTTCTTAGCATTTTTGACG* |
| 27 | GM72 | Diagnostic forward primers for *IPL1-HaloTag-TRP1* | *S. cerevisiae* genomic DNA | *GATAGAATGCGCCTTGGAGAC* |
| 28 | GM73 | Diagnostic reverse primer for *IPL1-HaloTag-TRP1* | *S. cerevisiae* genomic DNA | *CTGCGAATGCTCGTCTTGAG* |
| 29 | GM144 | Forward primer for *GLC7-HaloTag-TRP1* | pTSK573 | *AGCCAGCCCAAAAAAGTCTACCAAGGCAAGCTGGGGGTAGAAAGAAAAAACAAGCGGCCGCCGCTGCTGC* |
| 30 | GM145 | Reverse primer for *GLC7-HaloTag-TRP1* | pTSK573 | *TAATAAGTATTTTCCTTTTTAAACTTTGATTTAGGACGTGAATCTATTTACTATTTCTTAGCATTTTTGACG* |
| 31 | GM296 | Forward primer for *bub1*Δ | pAG32 | *GAAAGATTATTGACGGTTCCTATTGTTTGAATGTTAACGCTGACCAGGAAGCCAGCTGAAGCTTCGTACG* |
| 32 | GM297 | Reverse primer for *bub1*Δ | pAG32 | *GCAGGACACCAAAAAGTCACCTATGCGGGAGATGAAGGCATATTTATTCAGGCCACTAGTGGATCTG* |
| 33 | GM298 | Diagnostic forward primer for *bub1*Δ | *S. cerevisiae* genomic DNA | *GACGGTTCCTATTGTTTG* |
| 34 | GM299 | Diagnostic reverse primer for *bub1*Δ | *S. cerevisiae* genomic DNA | *GTGTTGTCATTGCTATGG* |
| 35 | GM1001 | Forward primer for *NDC10-3xGFP* tagging | pTSK405 | *GGCATGACCATCAAAATTCATTTGATGGTCTGTTAGTATATCTATCTAACCAAGCGGCCGCCGCTGCTGC* |
| 36 | GM1002 | Reverse primer for *NDC10-3xGFP* tagging | pTSK405 | *TACATGTCGGTATCCCTATACGAAACAGTTTAAACTTCGAAGCTCCCTCA GTTACTTGGTTCTGGCGAGG* |
| 37 | GM1003 | Forward primer for *IPL1-3xGFP* tagging | pTSK405 | *TGCATCCTTGGATACTAAGAAACAAGCCCTTTTGGGAAAATAAGCGGTTACAAGCGGCCGCCGCTGCTGC* |
| 38 | GM1004 | Reverse primer for *IPL1-3xGFP* tagging | pTSK405 | *GGGAGTGATTAATAGTGCCCTTCAAACGATTCTGTCATACTTTAATTCTAGTTACTTGGTTCTGGCGAGG* |
